# Fluorescent probes as markers of cell envelope structure and function in halophilic archaea

**DOI:** 10.64898/2026.02.20.706996

**Authors:** Elisa Ravaro, David J. Burr, Xavier Marques, Andreas Elsaesser, Adrienne Kish

## Abstract

Prokaryotes, particularly those in extreme environments, are capable of diverse metabolic states resulting in altered cell envelope structure and function. However, these changes are difficult to assess as standard fluorescent probes are often incompatible with extreme conditions and/or extremophile cell physiology. Halophilic archaea present the challenge of near-saturated intra-/extra-cellular salts, high membrane potential, and extended survival in altered metabolic states including entrapped within salt crystal fluid inclusions. We evaluated the compatibility of six fluorescent markers of cell envelope stability and activity with two model species, *Halobacterium salinarum* and *Haloferax volcanii*. Redox activity markers alamarBlue and pure resazurin solutions, membrane potential probes MitoTracker™ Orange-CMTMRos and Rhodamine 123, and SYTO 9 and propidium iodide (LIVE/DEAD™ kit) to assess cell membrane integrity were evaluated for use in bulk (microplate reader) and cell-specific (microscopy) applications. Limitations of each probe were identified, clarifying the utilization of each based on cell physiology, growth phase, medium composition, and probe exposure time including extended timescales needed to simulate the environmental conditions of haloarchaea. Of particular note, propidium iodide behavior was unreliable leading to double-labeling of cells and false interpretation of cells as dead. These data provide important insights into the study of prokaryotes in non-standard conditions.

## Introduction

Data on reduced metabolic states have forced reconsideration of classical conceptions of microbial dormancy (1, 2). This is particularly pertinent for microorganisms in extreme environments for which prolonged survival durations up to geological time scales have been suggested. Measuring such states, however, requires metabolic markers that function both in extreme physicochemical conditions and with the unique cell adaptations of extremophiles such as halophiles.

Halophilic archaea inhabiting hypersaline environments face fluctuating nutrient and oxygen availabilities, changes in salinity and brine composition, and variable energy sources. Studies of model species such as *Halobacterium salinarum* (*Hbt. salinarum*) and *Haloferax volcanii* (*Hfx. volcanii*) show that acclimation to changing conditions relies on well-regulated stress responses and metabolic versatility (see reviews (3, 4) and all references within). However, little is currently known about reduced metabolic states in haloarchaea, including dormancy. Metabolically inactive *Hfx volcanii* persister cells are genetically identical but phenotypically distinct, enabling survival under severe stress (5). Entrapment of haloarchaea, commonly *Halobacterium* (6), within halite fluid inclusions is an extreme stress condition that likely involves dormancy forms (7). Haloarchaea have been suggested to survive entrapment, in some cases based on observations of cell morphologies extracted from ancient geological formations (8). Precise ages of such samples remain controversial as currently only lithological and mineralogical dating methods are available (6). In addition, observed morphologies may represent active cells, structurally intact but dead, metabolically inactive or dormant states. The physiochemical properties of the brine itself can enable preservation of dead cells (9). The absence of validated *in situ* methods to study haloarchaeal cellular activity and structural stability has impeded testing these hypotheses.

The cell envelope is key to distinguishing between microbial activity states. In haloarchaea, a proteinaceous surface layer (‘S-layer’) cell wall maintains stability and is anchored into a diether isoprenoid lipid membrane (10) for generating redox gradients. The ‘salt-in’ osmotic strategy results in an usually high membrane potential for haloarchaea compared to other prokaryotes (11), providing a useful marker for metabolic activity.Novel dormancy states in low-nutrient, low-energy environments may involve cell envelope modifications (1), such as recycling of membrane lipids rather than *de novo* biosynthesis, which has been observed for halophilic archaea and evolutionarily-related methanogenic archaea (12) in energy-limited deep sediments. New tools are therefore needed to identify the metabolic state and structural stability of haloarchaeal cells, particularly in stress conditions.

One promising approach is the use of fluorescent probes targeting cell envelope membrane potential, redox-gradients or membrane structural integrity. However, to date few probes have been tested and verified with haloarchaea, often only compatible under specific applications. Membrane potential detection using the cationic tetramethylrosamine-based probe MitoTracker™ Orange-CMTMRos (MitoTracker) was validated in *Hbt. salinarum* strain S9 for epifluorescence microscopy applications (13). MitoTracker targets high membrane potential similar to that of mitochondria by binding covalent to protein SH-groups via its chloromethyl moiety, resulting in fluorescence quenching upon binding (14). It is retained by cells even after reduction in membrane potential, enabling labeling over several generations as observed in *Hbt. salinarum* (13). This study, however, did not evaluate MitoTracker-use with bulk fluorescence measurements such as platereader assays. Rhodamine 123, a lipophilic, cationic dye sensitive to both membrane potential changes and cellular ATP levels is an alternative probe for indicating metabolic activity or loss of metabolic function.

Redox activity is yet another method to differentiate active and inactive cells. Resazurin (blue color, abs=600nm, low fluorescence) is reduced by NADH or other biologically abundant reductive species in presence of cytoplasmatic reductases (15) to resorufin (pink color, abs=570nm, high fluorescence) which is measured by either colorimetric or fluorometric methods (16). Besides being widely used across bacterial and eukaryotic microorganisms to date compatibility of resazurin (either as pure resazurin sodium salt or commercial alamarBlue™ solutions) with haloarchaea has not been examined.

The few available fluorescent markers of cell membrane permeability are cytotoxic DNA intercalators including acridine orange, Hoechst 33342 (15) and the LIVE/DEAD™ kit based on the membrane-permeable SYTO 9 and membrane-impermeable propidium iodide (PI) (17). While the LIVE/DEAD kit has previously been used with haloarchaea (17), but studies reported discrepancies between colony-forming units (CFU) and PI-labeling (18) coherent with findings from bacteria (19). The underlying reasons for this discrepancy have not yet been identified, however recent studies in bacteria have shown that PI staining often yields false-positives due to high membrane potential and extracellular DNA (20), both of which are present in haloarchaea.

To use fluorometry to distinguish different metabolic and structural states of halophilic archaea further testing is necessary, specifically addressing discrepancies and identifying cell envelope-specific fluorescent probes compatible with haloarchaea. As such, the compatibilities of six fluorescent probes were tested, including two membrane-potential indicators (MitoTracker and Rhodamine 123), two resazurin-based indicators for redox-activity (resazurin sodium salt and alamarBlue™), and the LIVE/DEAD kit (SYTO 9 and PI). Compatibility was examined with *Hbt. salinarum* and *Hfx. volcanii* (10), two well-studied model species since different haloarchaea physiologies, reflective of their respective environments, could result in differential cell envelope behaviors. *Hbt. salinarum* is a hypersaline specialist (optimal growth at 4.28 M NaCl) with rod-shaped cells under optimal growth conditions. In contrast, *Hfx. volcanii* is a moderate halophile with smaller, irregularly disc-shaped cells, tolerant to a wide range of salt concentrations (1.8–3.5 M NaCl) (21) and higher concentrations of chaotropic Mg-salts.

The results of this study demonstrate both potential and limitations of using fluorescent probes to investigate cell integrity and activity in haloarchaea, using both bulk and cell-specific methodologies. The study parameters address the non-standard physicochemical and temporal conditions necessary to examine altered microbial cell states in extreme conditions These findings have broader implications for understanding similar processes in both archaea and bacteria.

## Results

Three complementary approaches were used to assess the cell envelope structure and function of fluorescently labeled haloarchaea: [i] bulk measurements with UV/Vis spectrophotometry to evaluate fluorescence signal development,compatibility and potential cytotoxicity of the fluorescent probes and their solvents with archaea in hypersaline conditions; [ii] cell-specific fluorescence microscopy for information on labeling efficiency and specificity with respect to both species and growth phase; [iii] compatibility of the fluorophores with extended incubation times to account for long-term analyses with haloarchaea under sub-optimal conditions such as within halite fluid inclusions where cell activity and division rates would be reduced. Epifluorescence microscopy was used to determine the labeling efficiency (ratio of labeled to unlabeled cells) and labeling specificity of the fluorescent probes with cells of both haloarchaea. In addition, heterogeneities resulting from changes in cell physiology were evaluated using exponential, stationary and decline growth phase cultures (two weeks after entering stationary phase).

### Redox probes

Compatibility of both resazurin formulations with different haloarchaea was evaluated using bulk measurements of relative fluorescence intensity (RFI) over time. Both alamarBlue and pure resazurin produced detectable fluorescent signals compared to growth medium blank controls (Fig. 1a, b). While the RFI of alamarBlue was similar for both haloarchaeal species tested, pure resazurin solution yielded 75% lower RFI in *Hbt. salinarum* (Fig. 1a) compared to *Hfx. volcanii* (Fig. 1b) after signal stabilization roughly 15 min following probe addition. This suggests that pure resazurin was more sensitive than the proprietary alamarBlue formulation to differences in either the physiology of the haloarchaeal species or the composition of their respective growth media (salinity, salt compositions, nutrient and organics sources). AlamarBlue therefore appears to be more suitable to a range of haloarchaea. Observations with epifluorescence microscopy revealed that the fluorescent signals did not localize to cells (Fig. 1c, d). High background fluorescence intensity was observed for all samples, likely due to the production of reduced resorufin by cells, which was released into the extracellular medium, resulting in magenta background on composite images (Fig 1c, d). This absence of cellular labeling is consistent with observations for mammalian cells, showing that resorufin does not accumulate intracellularly (22).

**Figure 1:**
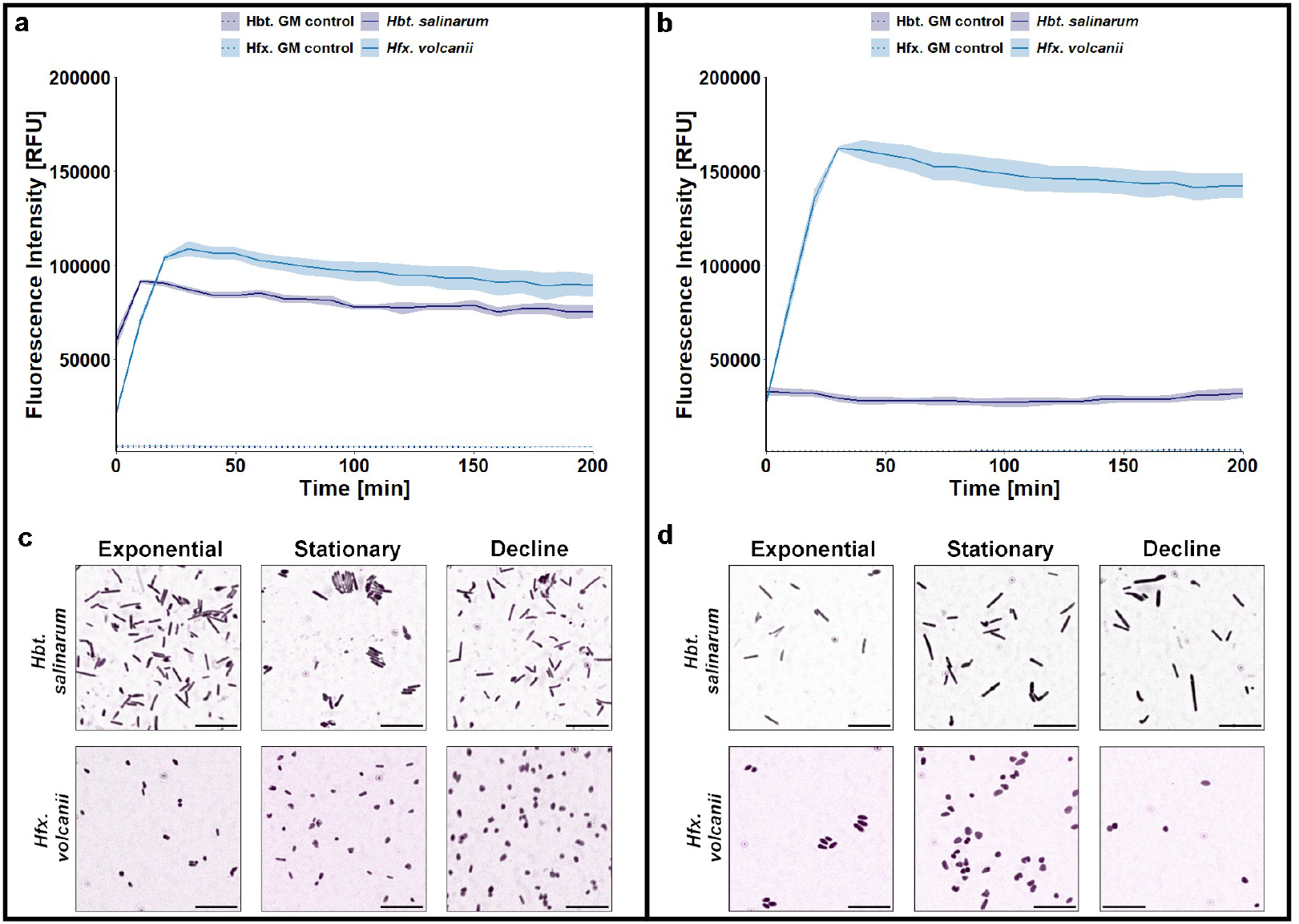
Compatibility of (a, c) alamarBlue and (b, d) pure resazurin salt with *Hbt. salinarum* and *Hfx. volcanii*. (a, b) Bulk measurements of relative fluorescence intensity over time were taken with cells of both organisms (stationary growth phase) and solely in growth medium (GM corresponding to CM+ and HvYPC respectively; dotted lines) at 37 ^°^C. Lines represent mean values, shaded regions represent the calculated standard deviation. Colors for alamarBlue samples (a, c) and pure resazurin samples (b, d) are identical since they represent the same detected molecule (fluorescent resorufin). Correlation of the bulk fluorescence signal with cells was analyzed with epifluorescence microscopy (c, d) for different growth phases (exponential, stationary and decline phases). Composite images that display the fluorescence signal in magenta and the cells in black, confirming that the fluorescent signals are not cell-specific. Scale bars indicate 10 µm. Stronger fluorescence intensity in *Hfx. volcanii* samples (a, b) is visible in microscopy images as magenta-filter, since fluorescent resorufin accumulates extracellularly, fitting to a higher relative fluorescence intensity in bulk measurements.

Since no cellular labeling was observed and given that resazurin can be reduced by non-cellular reductants such as ascorbic acid or reduced glutathione present in cell-free growth media (23), any artefactual effects with resazurin stemming from the growth media needed to be excluded. Reduction of alamarBlue was therefore measured at the maximal absorption wavelength for the oxidized molecule, resazurin (Abs600nm), after incubation in hypersaline solutions based on growth media of *Hbt. salinarum* and *Hfx. volcanii*. A decrease in absorption to 570 nm was expected if media components would reduce resazurin to resorufin. No reducing media components were identified for either media type (Fig. S1a), however different absorption intensities were observed for the solutions and alamarBlue™ (Fig. S1a). If non-neutralized peptone (PepNN) was used instead of recommended neutralized Peptone (PepN) the absorption of alamarBlue was lower, likely due to a more alkaline pH (Table S2). To account for potential degradative processes occurring with standard benchtop storage of haloarchaeal growth media, the same sterile media were tested again after 3 mo of storage (Fig. S1 b). All salt-based solutions with alamarBlue showed lower average absorption values after 3 mo benchtop storage. These observations indicate that it is best to use freshly prepared growth media with any resazurin-based redox probes, especially if solutions are non-buffered or have more alkaline pH. After reducing effects of the hypersaline growth media were excluded, the resazurin-based probes were tested for cytotoxicity in haloarchaea. AlamarBlue is marketed as non-toxic probe, if used according to the manufacturer-recommended incubation times. However, incubation with resazurin over the course of microbial growth exceeds this time. Therefore, the potential toxicity of resazurin-based probes was tested with both haloarchaea over 60 hours. Growth inhibition of *Hbt. salinarum* and *Hfx. volcanii* was observed with both probes (Fig. S2). This corresponds to findings (24) that caution users of cell-type dependent levels of resazurin-tolerance. Since resazurin (alamarBlue™) is known to be reduced by electron carriers such as NADPH/H+ that are essential to maintaining redox gradients for ATP production, a depletion of these carrier-molecules might inhibit the cellular respiratory chain (24).

### Membrane potential probes

Bulk measurements, were performed since literature currently does not describe measurement of RFI of the described membrane potential probes with haloarchaea. Measurements with both probes showed varied signal to noise properties relative to both probe type and haloarchaeal species. Stationary phase *Hfx. volcanii* and *Hbt. salinarum* cells incubated with MitoTracker produced relatively stable fluorescence signals over time (Fig. 2a). However, MitoTracker also produced fluorescent signals in cell-free controls. Fluorescence in *Haloferax* growth medium (HvYPC) stabilized after 20 minutes whereas in *Halobacterium* growth medium (CM+) it increased over time. Additionally, cell-specific fluorescence intensities were 1.4-fold higher in labeled *Hfx. volcanii* compared to *Hbt. salinarum*. These observations complicate the interpretation of bulk measurements for *Hbt. salinarum*, where fluorescence was evaluated based on its dominant quenching effect and prolonged incubation times (>2 h). Under these conditions, MitoTracker RFI varied depending on both salinity and organism type.

**Figure 2:**
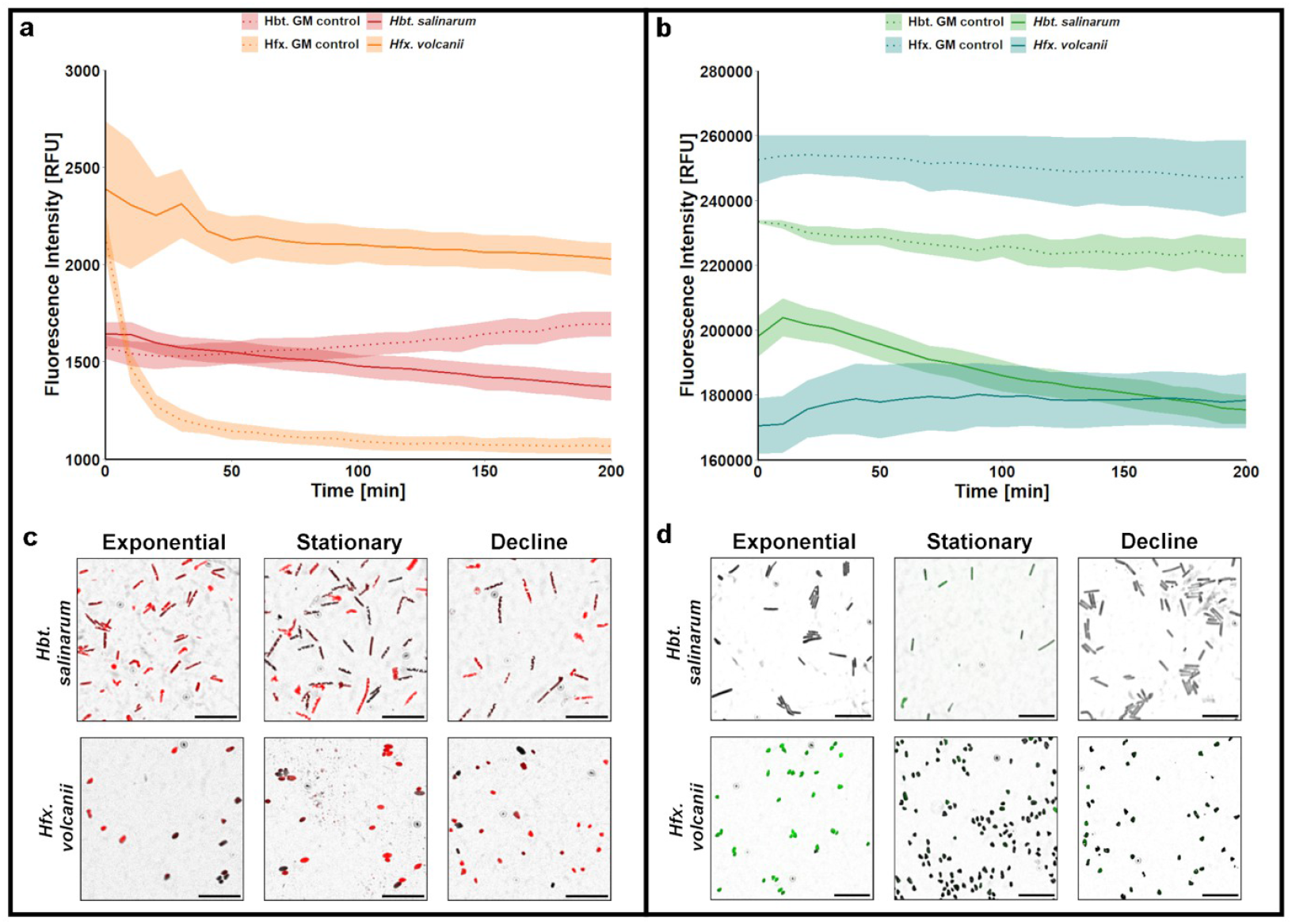
Compatibility of (a, c) MitoTracker orange CMTMRos and (b, d) Rhodamine 123 with *Hbt. salinarum* and *Hfx. volcanii*. (a, b) Bulk measurements of relative fluorescence intensity over time were taken with cultures of both organisms as well as with growth medium controls (GM=growth medium, corresponding to CM+ for *Hbt. salinarum* and HvYPC for *Hfx. volcanii*; dotted line) at 37^°^C. Mean values as lines/dots, calculated standard deviation as shaded regions. Correlation of the bulk fluorescence signal with cells was analyzed with epifluorescence microscopy (c, d) for different growth phases (exponential, stationary and decline). Composite images that display the fluorescence signal in red (MitoTracker) or green (Rhodamine 123) and the cells in black. Scale bars indicate 10 µm. Cell specific labeling could not be observed for Rhodamine 123, where only few cells showed weak fluorescent signals.

Rh123 showed a stronger quenching effect than MitoTracker in haloarchaea, with cell-specific fluorescent signals being ∼1.3-fold lower compared to Rh123 in cell-free controls (Fig. 2b). Similar quenching of Rh123 has been reported in bacteria (25) due to high probe concentrations, insufficient washing, or high salinity, particularly elevated K+ concentrations (26). The concentrations used here fall within the previously determined range for *Hfx. volcanii* (27). Notably,cell-free medium controls revealed higher Rh123-fluorescence in HvYPC (*Hfx. volcanii* medium) than in CM (*Hbt. salinarum medium*), coherent with the >2-fold higher K+ concentration of HvYPC.

Epifluorescence microscopy was used to determine if growth phase specific differences in cellular labeling and therefore membrane-potential occur in either *Hbt. salinarum* or *Hfx. volcanii*. Non-uniform labeling of both species was observed for both membrane potential probes (Fig. 2c, d). These results were verified by enumeration, showing that the proportion of MitoTracker labeled cells was growth phase independent, remaining between 80-90 %, with *Hfx. volcanii* showing slightly lower average numbers of MitoTracker-labeled cells than *Hbt. salinarum* (Fig. S3). Incomplete labeling with MitoTracker is consistent with the findings of Maslov et al. (13) for *Halobacterium*, though direct comparisons are limited due to differences in microscopy techniques used between the two studies (confocal vs. epifluorescence microscopy). The observed heterogeneity may reflect cell-to-cell variations in membrane potential (ion channel density, permeability, ATP levels) or occur due to different probe accumulation inside the cells. Protocol optimization (for example, 1x washing instead of 3x before microscopy, incubation-times >100 min) enhanced signal discrimination. In contrast Rh123 produced low fluorescence intensities (Fig. 2d), resulting in a labeling efficiency below 40 % (Fig. S3). Either the strong probe quenching observed in bulk measurements (Fig. 2b), rendered the background fluorescence indistinguishable from labeled cells, or Rh123 might have been lost from haloarchaeal cells during the manufacturer-recommended wash steps intended to eliminate such background signal interference. While both MitoTracker and Rh123 are lipophilic cationic probes, differences in side-groups result in MitoTracker being retained in cells even after a loss of membrane potential, while Rh123 is lost from cells, which may result in loss of Rh123 probes during wash steps. Additionally, *Hfx. volcanii* cells export Rh123 via an ATP-driven transporter in the presence of an excess of amino acids (28). Any effect on signal detection derived from either the probe concentrations or the use of generic filter sets simply reinforces the requirement of Rh123 for a high degree of preliminary optimization if it is to be used with haloarchaea.

Neither MitoTracker or Rh123 exhibited cytotoxicity in *Hbt. salinarum*, supporting findings of Maslov et al. (13) for MitoTracker. The presence of MitoTracker conveyed a slight growth advantage during the late exponential growth phase, whereas Rh123 caused a slight decrease in the mid-exponential to early stationary growth of *Hbt. salinarum* (Fig. S2a). The DMSO solvent used for both probes resulted in a concentration-dependent growth inhibition of *Hbt. salinarum* at concentrations over 2.5 % (Fig. S4) with total inhibition occurring at 10 %, consistent with previous findings for bacteria (29). This highlights the importance of stock solutions concentration (Table S1) to prevent DMSO cytotoxicity after addition to growth media. Concentrations of <1 % have a cell-preserving effect preventing passage from stationary to decline phase. As *Hbt. salinarum* can use low concentrations of DMSO for anaerobic growth (30), it is possible that cells underwent a metabolic switch that resulted in the observed prolonged survival in a reduced metabolic state. Therefore, attention should be paid to the DMSO concentration used with metabolic markers.

### LIVE/DEAD Viability/Cytotoxicity kit

Prior to testing the LIVE/DEAD kit compatibility with haloarchaea, *Escherichia coli* cultures in stationary growth phase were used as a reference to validate kit functionality and establish a baseline for comparison. CLSM confirmed uniform labeling of live *E. coli* cells with SYTO 9, and dead cells (permeable cell envelopes) with PI (Fig. 3a). Quantification of labeling efficiency and specificity (Table 1) showed only 2 % of *E. coli* cells remained unlabeled and that PI efficiently replaced SYTO 9 in dead cells as expected. In comparison, *Hbt. salinarum*, and to a lesser degree *Hfx. volcanii*, showed higher variability (standard deviation) in labeling efficiency in stationary phase, where both live and dead cells are expected. This is reflective of the polyploidy state of each species, and high mutation rates in haloarchaea (31–34).

**Table 1:** Effect of incubation conditions with the LIVE/DEAD Viability/Cytotoxicity kit on cell labelling of *E. coli, Hbt. salinarum* and *Hfx. volcanii* (n=3 for each, with a minimum of 100 cells counted per biological replicate). Stationary phase cultures were either labelled immediately while decline phase cultures were stored for 5 d on the benchtop with labelling either at the end of the storage period (post-storage) or throughout the entire 5 d (intra-storage). The numbers of total, single and double labelled cells were counted and percentage calculated as number of fluorescent cells divided by total cell counts in brightfield, presented as an average across replicates. High standard deviations in particular for *Hbt. salinarum* are likely due to the genetic variability of halophilic archaea.

| Species | Labeling protocol | total labelled cells [%] | single SYTO 9 labelled cells [%] | single PI labelled cells [%] | double labelled cells [%] |
| --- | --- | --- | --- | --- | --- |
| *E. coli* | Stationary | 98 ± 0 | 95 ± 0 | 5 ± 0 | 0 ± 0 |
| *Hbt. salinarum* | Stationary | 87 ± 24 | 85 ± 24 | 5 ± 5 | 9 ± 19 |
|  | Post-storage | 74 ± 26 | 64 ± 17 | 7 ± 7 | 29 ± 17 |
|  | Intra-storage | 44 ± 26 | 28 ± 36 | 53 ± 37 | 19 ± 19 |
| *Hfx. volcanii* | Stationary | 95 ± 8 | 1 ± 2 | 2 ± 16 | 97 ± 16 |
|  | Post-storage | 5 ± 15 | 83 ± 11 | 4 ± 2 | 13 ± 9 |
|  | Intra-storage | 94 ± 7 | 1 ± 2 | 49 ± 16 | 5 ± 16 |

**Figure 3:**
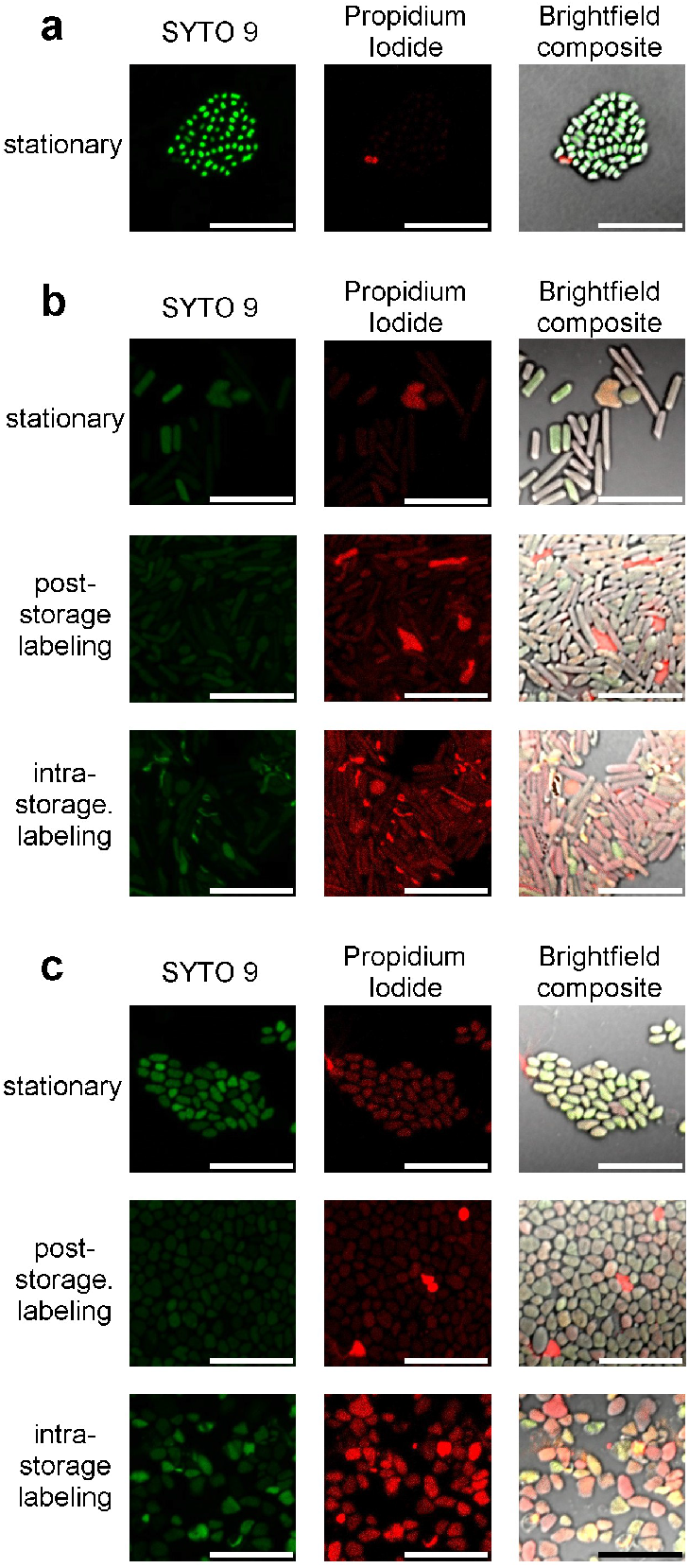
CLSM of various organisms incubated with the LIVE/DEAD™ BacLight Viability/Cytotoxicity kit (L/D). (a) *E. coli* labelled in stationary growth phase with L/D. (b) *Hbt. salinarum* and (c) *Hfx. volcanii* in stationary growth phase, labeled after 5 days of incubation (post-storage labeling) or incubated for 5 days with L/D (intra-storage labeling). Scale bars indicate 10 µm. Double-labeling of cells is depicted as yellow-orange color in brightfield composite images.

Interspecies variability in SYTO 9 fluorescence was confirmed by bulk measurements, showing a 2-fold higher fluorescence intensity in *Hfx. volcanii* than in *Hbt. salinarum* (Fig. S8), while PI did not produce a detectable fluorescence signal distinguishable from the growth medium control in either species, suggesting a role for salinity and/or haloarchaeal physiology in fluorescence intensity. Importantly, haloarchaeal cells displayed double-labeling with overlapping SYTO 9 and PI signals producing yellow-orange cells in composite images (Fig. 3b, c brightfield composite panels). This contradicts the expected probe behavior for dead cells as PI should displace SYTO 9 due to its greater DNA binding affinity (19). Additionally, any remaining SYTO 9 signal should be locally absorbed within a ∼10 nm range by PI due to Förster resonance energy transfer (FRET) (35) as FRET-efficiency is inversely proportional to distance between energy donor and acceptor. Given that PI is assumed to be membrane-impermeable, a double label signal localized to a cell would therefore require either a live cell with PI bound to extracellular DNA in close proximity to the cell surface, or PI entering live polyploid cells at a slower rate creating a mosaic effect. This identifies PI as the source of the inaccuracies for use of the LIVE/DEAD kit to determine survival in haloarchaeal cultures.

To determine the extent of this phenomenon labeling specificity was calculated as single-labeled cells in proportion to the total number of labeled cells. *E. coli* cells in stationary phase showed no double-labeling, whereas the phenomenon was particularly prevalent in *Hfx. volcanii* with 97 % of stationary-phase cells displaying both fluorescent signals (Table 1), annulling any interpretation of cell viability.

Growth phase had a significant and species-specific impact on labeling efficiency. Stationary phase cultures were compared to those stored at RT for an additional 5 days prior to labeling (‘post-storage’). The data suggests that physiological changes of *Hfx. volcanii* cells during storage after stationary phase produced over 7-fold fewer labeled cells, but those labeled had a higher specificity for either SYTO 9 or PI evidenced by a 2-fold reduction in double-labeled cells (Table 1). Conversely, comparable numbers of *Hbt. salinarum* cells were labeled in both stationary and post-storage, but with a 3-fold increase in doubling labeling.

To further investigate the potential for labeling haloarchaeal cells in reduced metabolic states as within halite fluid inclusions or during routine benchtop storage of cultures, the labeling time was extended to the 5-day storage period (‘intra-storage’ labeling). This is an atypical application of the LIVE/DEAD kit as both SYTO 9 and PI are DNA intercalating probes, assumed to result in death of haploid cells during cell division. However, previous studies have successfully applied this method to polyploid haloarchaea in halite (36), which suggests that survival of such cells is either a temporary phenomenon if indeed entrapped cells no longer undergo cell division, or the product of labeling only a sub-population of genome copies per polyploid cell, or false-positive signals. This extended labeling time resulted in a high heterogeneity for *Hbt. salinarum*, with a nearly 2-fold reduction in total labeled cells (Table 1, inter-vs. post-storage). A corresponding reduction of double labeling under these conditions was indicative of higher specificity, with no visible changes in cell morphology evident in brightfield composite images (Fig. 3b).

Validating the interpretation of PI-labeled cells as dead is needed as membrane permeability in haloarchaea may not necessarily represent cell death, contributing to double-labeling. Standard ethanol treatment of un-fixed cells in both strains used here resulted in cell lysis rather than membrane permeability (see Supplementary Methods) and no method has been found yet to produce a dead-but-intact (non-permeabilized) haloarchaeal cell. An alternative is CFU counts. As the DNA-intercalating action of both PI and SYTO 9 leads to inhibited cell division, CFU represent cells surviving LIVE/DEAD labeling (at least one genome copy remaining unblocked by DNA intercalators) or the respective storage condition. This method showed that labeling of stationary cells did not result in lower CFU compared to unlabeled cells in the same growth phase (N/No) while *E. coli* was susceptible to the toxicity of LIVE/DEAD (Fig. S5). Post-storage and intra-storage labeling however did result in a 50 % reduction of *Hbt. salinarum* CFU in comparison to an unlabeled storage control (Fig. S5, storage control), indicating a physiological change that made *Hbt. salinarum* more susceptible towards the toxicity of DNA-intercalating probes, possibly allowed by its decreasing polyploidy (31– 34). However, an increase in CFU during the storage period of unlabeled *Hbt. salinarum* prevented any conclusions about actual dead-cell numbers in the samples analyzed by microscopy. In contrast a CFU reduction of 55 % could be shown for *Hfx. volcanii* between the storage control and stationary samples (Fig. S5). Since CLSM dead cell counts (Table 1; single-labeled PI cells) indicated only 4 % of post storage labeled *Hfx. volcanii* cells as dead, a difference of 50 % in estimated numbers of dead cells emerges between these two methods. In addition, this method validated anecdotal evidence that *Hbt. salinarum* survives better and in fact continues dividing during benchtop storage compared to *Hfx. volcanii*.

## Discussion

Hypotheses of extended longevity and novel dormancy modalities in prokaryotes require new methods of investigation. This is of particular importance for extremophilic microorganisms. The effects of both the cell envelope adaptations of extremophiles and the physiochemical conditions required for their growth must be taken into consideration when measuring cell integrity and activity. While this study focused on the compatibility of fluorescent markers for redox activity, membrane potential, and cell membrane integrity with haloarchaea, the results also revealed some principles widely applicable to other prokaryotes.

The feasibility of resazurin-based bulk redox activity measurements in halophilic archaea was validated, if both the chemistry of the fluorescent probe and the precise composition of the hypersaline growth medium are considered. Since fluorescence signals originate mainly from extracellularly accumulated resorufin, the probes are not recommended for cell-specific analyses like flow-cytometry or fluorescence microscopy. Long-term incubation is not recommended due to probe-toxicity.

For membrane potential, the results support the use of MitoTracker as a probe for bulk RFI measurements in haloarchaea, conditioned on preliminary compatibility testing with both the strain and suspension medium. Attention must be paid to quenching by selecting appropriate controls for interpreting fluorescence signals in bulk spectrometry. When the application protocol is optimized (1x washing instead of 3x before microscopy, incubation-times >100 min), MitoTracker is the most versatile for use under a variety of conditions and different halophilic organisms. However, the exhibition of uneven fluorescence requires further investigation to determine whether these differences reflect genuine variations in membrane potential or are influenced by other physiological factors. Furthermore, the overall weak fluorescence intensity of MitoTracker and its permanent binding to cellular structures (14) prevents the monitoring of dynamic changes in membrane potential, highlighting the need for further research to identify more suitable probes for assessing membrane potential fluctuations. While Rh123 has previously been used with *Hfx. volcanii* in flow-cytometry (28), amino acid concentrations in the growth medium affect retention. The data here provide additional restrictions on Rh123 application emphasizing strict optimization of experimental conditions. The strong background fluorescence of Rhodamine 123 and its potential incompatibility with washing steps necessary to increase signal-to-noise ratios suggest that this membrane potential indicator should only be applied after species-specific protocol optimization. Fluorescence bulk measurements showed that MitoTracker and Rh123 should be evaluated upon compatibility on a case-by-case basis, including both physicochemical conditions such as salinity as well as the microorganism itself.

Evaluation of the LIVE/DEAD kit to study cell membrane integrity over extended duration and in extreme conditions raised important concerns for its use, challenging the reliability of this type of labeling even under controlled laboratory conditions. High variability in labeling efficiency and specificity between the two tested haloarchaea species and E. coli highlights a strong species bias for the LIVE/DEAD kit within the prokaryotes. Even within haloarchaea, species bias was evidenced by differences between the two species in the labeling efficiency across various growth phases. The changes observed after culture storage also present a significant limitation for applications with haloarchaea as cultures are routinely stored in laboratories at room temperature after reaching stationary growth phase. In addition, the storage condition is also reflective of the hypothesized physiological state of *Halobacterium* cells entombed within the fluid inclusions of halite (7). Taken together, these high variabilities rendered data interpretation with the LIVE/DEAD kit qualitative at best.

More importantly, PI failed to displace SYTO 9 consistently across different species, giving “false-dead” or double-labeled signals. The reliability of PI with some model organisms like *E. coli* resulted in its widespread application including with haloarchaea such as *Hbt. salinarum* (17, 37, 38), but inconsistencies between viable cell enumeration and CFU plate counts (17) have been previously noted under certain conditions. Contributing factors, such as the high-salt environment, the presence of extracellular DNA, and particularly the elevated membrane potential in haloarchaea, result in this false-positive PI labeling.

These results have implications far beyond halophiles by contributing to a growing body of literature for prokaryotes showing the limitations of PI for detection of dead cells. The observed differences in SYTO 9 and PI labeling are particularly restrictive for automated analyses such as flow-cytometry, independent of cell type. Furthermore, users should maintain set incubation times since extended exposure of cells to PI and SYTO 9 alters results, impacting reliability for batch analytics. Based on these findings, use of PI is not recommended as a dead-cell marker for mixed communities, such as environmental samples, or for single organisms that have not yet an optimized usage protocol (such as *Hfx. volcanii*).

Taken together, this study provides new insights into evaluating cell envelope integrity and activity in extremophiles using both cell-specific and bulk methods. These findings open new doors to discover novel prokaryotic dormancy and cell preservation modes in extreme conditions.

## Material and Methods

All experiments were performed as biological triplicates, with additional technical replicates as indicated.

### Culture conditions

*Haloferax volcanii* (DSM3757) was routinely grown under oxic conditions in HvYPC complex medium (see Table S1). For solid media 1.5 % agar (w/v) was used. After autoclaving 3 mM CaCl_2_ was added.

*Halobacterium salinarum* strain NRC-1 was grown under oxic conditions in either complex media (CM) (see Table S1) or complex medium plus (CM+, derived from C. Evilia pers. comm) additionally containing 0.5 % (v/v) glycerol, and 0.22 µm filter sterilized solutions of trace metals that were added after autoclaving. For solid media 2 % agar (w/v) was added.

Liquid cultures were incubated at 37 ^°^C, 180 rpm in glassware that had been vigorously rinsed multiple times in MilliQ® water after washing and prior to sterilization to remove all traces of detergents that have detrimental effects on the cell envelopes of haloarchaea. Cultures were grown to an optical density at 600 nm (OD600nm) of 0.5 (exponential growth phase), 1.0-1.5 (stationary growth phase) or incubated for a further 2 weeks after reaching stationary growth phase (corresponding to decline phase).

For LIVE/DEAD experiments, attempts were made to produce cells with permeabilized membranes as a control for propidium iodide labelling using a standard 60 % ethanol treatment with both *Hfx. volcanii* and *Hbt. salinarum*. Both these strains of noncoccoid haloarchaea were not compatible with ethanol permeabilization resulting in cell lysis, coherent with the results from Leuko *et al*. 2004 (17). Survival was instead enumerated for live cells by CFUs grown on nutrient agar. Inoculated plates were dried in a laminar flow hood prior to incubation at 37 ^°^C inverted inside sealed bags in the presence of a moistened paper towel forming a humidity chamber to avoid salt crystallization for 6-7 days. Viability was expressed as a percent % survival compared to control cultures (N/N0 × 100). Statistical analysis was performed using two-tailed unpaired equal and unequal variance students t-tests, where p < 0.05 was considered significant.

Statistical analyses for CFU counts were performed using two-tailed unpaired equal and unequal variance students t-tests, where p < 0.05 was considered significant.

### Fluorescent probe preparations and cell labelling

Stock solutions of fluorescent probes were prepared according to manufacturers’ protocols with either DMSO or MilliQ® water as solvent (Table S2). Unless otherwise stated, all samples were incubated in the dark on a tube revolver/rotator (ThermoScientific® reference 88881002, 20 rpm) according to the labeling protocol presented in Table S2. Where wash steps are indicated, the sample was centrifuged 3000 x G, 3 min, room temperature (RT), the supernatant discarded and the cell pellet resuspended in growth medium.

PI and SYTO 9 of the LIVE/DEAD™ BacLight™ Bacterial Viability Kit (Invitrogen™ reference L7012) were incubated according to Table S2 for immediate imaging. Alternatively, for the ‘intra-storage’ labeling condition these fluorescent probes were kept (in the dark, RT, without agitation) in growth medium with haloarchaeal cells for 5 days and then imaged.

### Bulk measurements

All bulk measurements were performed using a POLARstar® Omega (BMG Labtech) spectrophotometer running the Omega software (version 5.70 R2). Growth curves were measured in absorbance mode (Abs600nm), as was cross-reactivity of resazurin in deconstructed high-salt complex media for *Hfx. volcanii* and *Hbt. salinarum* (see Supplementary Methods). The shift in absorption was followed from the oxidized, blue resazurin (600 nm) to the reduced, pink resorufin (570 nm).

Bulk measurements of relative fluorescence intensity (RFI) were performed in fluorescence mode using cultures in stationary growth phase analyzed as technical duplicates of each biological replicate and mixed with the fluorescent probes (Table S1).

Data of absorbance- and RFI-bulk measurements was background corrected by either blank controls of cells in growth medium or pure growth medium.

### Microplate growth measurement

Growth curves were measured in transparent Nunc™ MicroWell™ 96-Well DeltaSurface plates (Thermo Scientific™) with a sample volume of 200 µl as technical duplicates per biological replicate (3 x 2 = 6 replicates total). Cultures were diluted to OD600nm=0.05 with growth medium, mixed with the fluorophores at the indicated final concentrations and incubated over 70 hours, 37 ^°^C, 200 rpm, double-orbital shaking in a POLARstar® Omega (BMG Labtech) spectrophotometer running the Omega software (version 5.70 R2) with (growth medium blank-corrected) OD600nm measured every 15 min for the first 4 hours and then every hour. To prevent evaporation, empty wells were filled with water and the plates sealed with parafilm. Pure DMSO (Invitrogen™ reference D12345) was incubated with *Hbt. salinarum* at increasing concentrations (0.1, 0.2, 1, 2.5, 5 and 10 % (v/v)) under the same conditions to test the growth impact of this commonly used solvent.

### Fluorescence intensity bulk measurement

Bulk measurements of relative fluorescence intensity (RFI) were performed in black bottom Nunc™ MicroWell™ 96-well plates (Thermos Scientific™, reference 137101) with a sample volume of 200 µl. Cultures in stationary growth phase were analyzed as technical duplicates of each biological replicate and mixed with the fluorescent probes in a final volume of 1 mL (Table S2). Samples were directly added to the wells and incubated for up to 4 h at 37 ^°^C and 200 rpm in a POLARstar® Omega platereader in fluorescence mode. Measurements were performed with the appropriate fluorophore specific filter sets (Table S2). RFI of bulk measurements with cells were determined for each fluorescent probe by subtracting the raw value (measured as relative fluorescence units, or RFU) for cells in growth medium without the fluorescent probe from value of the equivalent sample with the fluorescent probe. For autofluorescence controls, RFU of pure growth medium was subtracted from RFU of growth medium with the fluorescent probe.

### Resazurin reduction experiment

Cross-reactivity of resazurin was tested in deconstructed high-salt complex media for *Hfx. volcanii* and *Hbt. salinarum*. Solutions contained either only the salts needed for each organism (HfxS and HbtS) or the salts plus only one of the main complex organic nutrient sources (Table S1): Oxoid® neutralized peptone (Pep N), Oxoid® non-neutralized peptone (Oxoid® LP0037; Pep NN) Bacto™ yeast extract (gibco™ 12750; YE) and Bacto™ Casamino acids (gibco™ 223050; Cas). Test solutions were mixed with alamarBlue™ as indicated above (six technical replicates per sample) and 200 μl of each sample were added into transparent Nunc™ MicroWell™ 96-Well DeltaSurface plates (Thermo Scientific™). Absorption of the oxidized, blue resazurin, at 600 nm (Abs600nm) was measured using a POLARstar® Omega spectrophotometer at 37 ^°^C for 200 min and 200 rpm with meander corner well shaking and additional 300 rpm double orbital shaking every 15 min. In case of resazurin-reduction to resorufin, the absorbance shifts to 570 nm (pink resorufin).

### Cells-specific observations using epifluorescence/confocal microscopy

Epifluorescence microscopy was performed on cells immobilized on agarose gel slices using a Nikon TE300 inverted microscope, imaged with a 100x oil immersion objective(NA 1.3) in brightfield (phase contrast 3) and with fluorescence filter sets compatible with each fluorophore (Table S1). Images were acquired using a Photometrics CoolSNAP HQ camera and the MetaMorph® software. Confocal Laser Scanning Microscopy (CLSM) of cells labelled using the LIVE/DEAD kit was carried out using an inverted LSM 880 Zeiss microscope equipped with a 40x oil immersion objective (NA 1.3) using the Zeiss Zen blue software. Images were analyzed using the FIJI software (Fiji is just ImageJ; (39)). CLSM images were acquired as Z-stacks and therefore a Z-projection of median intensities for brightfield images and of maximal intensity for fluorescence images was analyzed. Single epifluorescence images were analyzed, and background of all images was subtracted by a rolling ball radius of 20 pxl. Brightness and contrast were adjusted (for both CLSM and widefield images). Composites were generated by merging a fluorescent filter image with the respective brightfield image, displaying the brightfield in grey and the fluorescent image in either magenta, green or red. CLSM images of the LIVE/DEAD™ labeled cells were counted to calculate labeling efficiency and specificity of each image. Visible cells in areas of 200×200 pixels (20×20 µm) per image (separate for each image in brightfield, SYTO 9-filter, PI-filter) were counted manually. Fluorescence composites (PI- and SYTO 9 composite) were used to identify number of double-labeled cells (yellow, orange and dark orange colors). Three different labeling protocols were used to simulate potential applications ranging from optimal to suboptimal environments for haloarchaea in stationary growth phase, such as entombment in halite fluid inclusions. These included: (i) the standard labeling protocol as used for *E. coli* for stationary phase cultures (all cultures grown at 37 ^°^C, 180 rpm), (ii) continuous exposure to SYTO 9 and PI throughout a secondary 5-day benchtop incubation (no shaking) after reaching stationary phase (intra-storage labeling), and (iii) labeling after the secondary 5-day incubation (post-storage labeling) compared to unlabeled storage control samples. While the latter two protocols followed the passage of cultures from stationary to decline growth phase, intra-storage labeling began in stationary phase whereas post-storage labeling only involved decline phase cells. Labeled cells were manually counted and labeling efficiency was calculated as total labeled cells (%) in proportion to the total number of visible cells in brightfield.

## Acknowledgements

The authors acknowledge the support of the French National Research Agency (ANR), under grant ANR-21-CE49-0017 (project ExocubeHALO), CNES programme Exobiologie focused on the Exocube space experiment. ER was supported by PhD grants from CNES programme Exobiologie (project Fleur de Sel) and the ANR as part of the France 2030 initiative (project CRUSTAL). The authors thank Cyril Willig for assistance in use of the CeMIM (MNHN) microscopy platform. For the helpful discussions and share of ideas the authors gratefully acknowledge Lucas Bourmancé, Ruben Nitsche, Antonio Ricco, Sergio Santa Maria, Samuel Marre, and Anaïs Cario.

## Authors’ Contribution Statement

ER designed and performed the experiments, analyzed the data and wrote the draft manuscript; XM assisted with microscopy observations and data analyses; DJB and AE participated in designing fluorophore experiments and data interpretation; AK provided the project funding, designed and supervised the experiments, participated in data analyses and manuscript preparation. All co-authors contributed to revising the manuscript.

## Supplementary information

Online version contains supplementary materials (Fig. S1, S2, S3, S4, S5, S5.1, Tab. S1, S2).

