## Supplementary Material for "Fluorescent probes as markers of cell envelope structure and function in halophilic archaea"

### Supplementary Figures

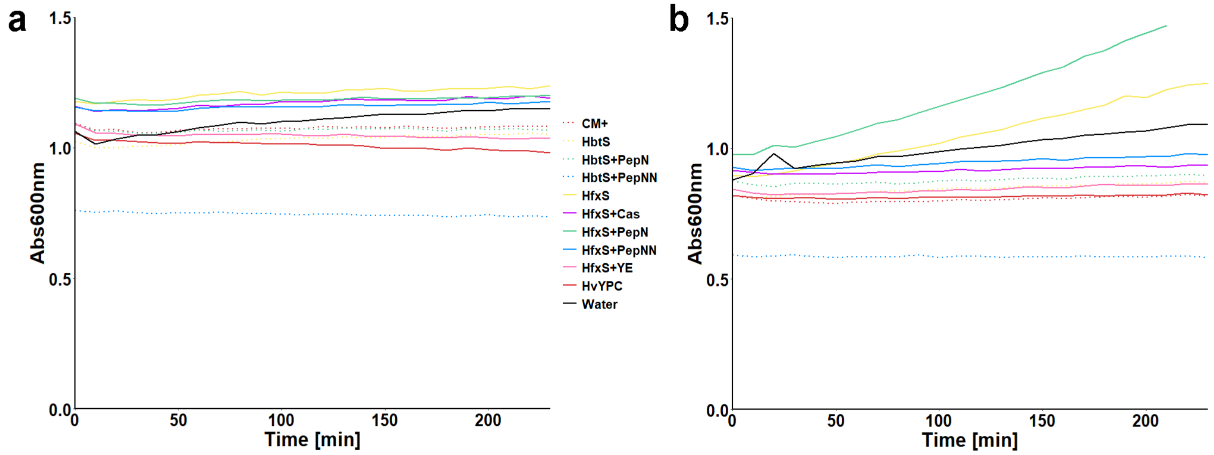


**Figure S1:** Effects of high-salt growth media composition on bulk measurements of alamarBlue™. Cell-free solutions were prepared representing complex growth media (HvYPC for *Hfx. volcanii* and CM+ for *Hbt. salinarum*) as well as deconstructed media solutions: basal salt solutions without organics (HfxS and HbtS), basal salts with non-neutralized peptone (HfxS/HbtS+PepNN), and basal salts with neutralized peptone (HfxS/HbtS+PepN). For Hfx. volcanii only, basal salts with yeast extract (HfxS+YE) or with casamino acids (HfxS+Cas) (see Table S2 for compositions of each solution). Each high-salt solution was prepared, filter sterilized and incubated with the respective resazurin-based probe in a transparent 96 well-plate at 37 °C, 200 rpm in POLARstar® Omega spectrophotometer. Absorbance at 600nm (Abs600nm) of each cell-free solution was measured over time to identify formation of resorufin (pink color, maximal absorption at 570nm) following the reduction of resazurin (blue color, maximal absorption at 600 nm). Freshly made solutions were first incubated with alamarBlue™ and the Abs600nm measured (panel A), showing a decreased reduction of resazurin with respect to pH, based on the lower Abs600nm values in the presence of non-neutralized peptone (PepNN). A second set of solutions were then stored for 3 mo at RT and incubated with alamarBlue™ (panel B) to account for potential degradative processes occurring during extended storage periods for growth media. All salt-based solutions with alamarBlue showed lower absorption values after 3 mo benchtop storage, indicating a strong effect for the age of growth media on bulk measurements of haloarchaeal redox activity. Results indicate that it is best to use freshly prepared growth media with any alamarBlue™, especially if solutions are non-buffered or have more alkaline pH.

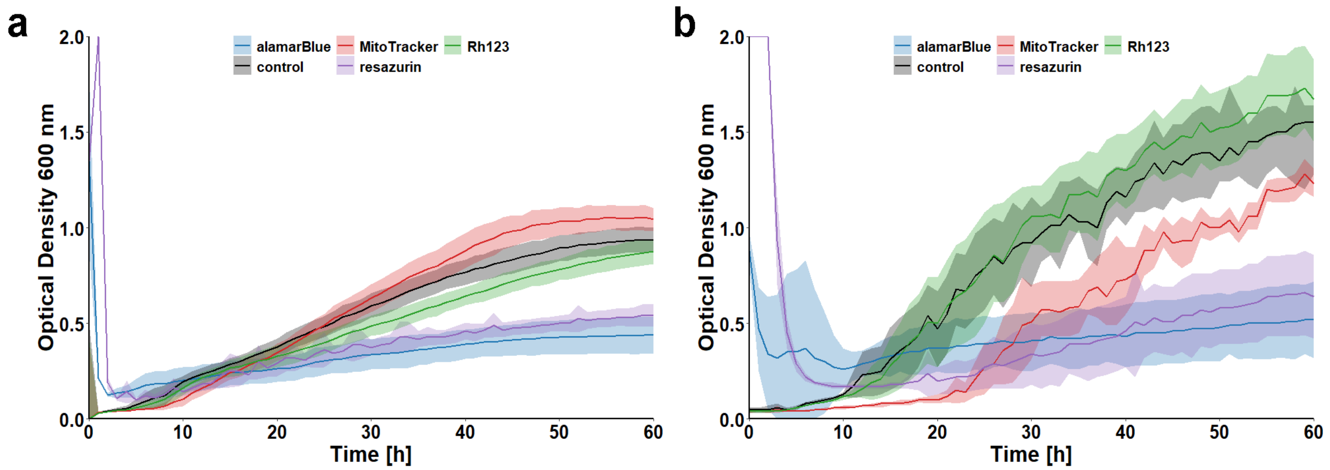


**Figure S2:** Growth impact of fluorescent probes on (A) *Hbt. salinarum* and (B) *Hfx. volcanii*. Organisms were grown in complex medium with alamarBlue™, resazurin, MitoTracker™ Orange CMTMRos, Rhodamine 123 (Rh123) and without fluorophores (control) in 96-well plates to measure OD600nm over time (37 °C, 200 rpm, double-orbital shaking in a POLARstar® Omega (BMG Labtech) spectrophotometer running the Omega software (version 5.70 R2) with blank-corrected OD600nm measured every 15 min for the first 4 hours and then every hour). The initial increase in OD600nm in the presence of alamarBlue™ and resazurin for both species (time: 1 to 4h) was due to absorption overlap at 600 nm of blue resazurin (prior to its reduction by redox-active cells to the pink resorufin) and haloarchaeal cells. Data over this timecourse show that resazurin-based probes (alamarBlue and pure resazurin) inhibited growth of both organisms, whereas no cytotoxicity was displayed in the presence of either MitoTracker™ or Rh123.

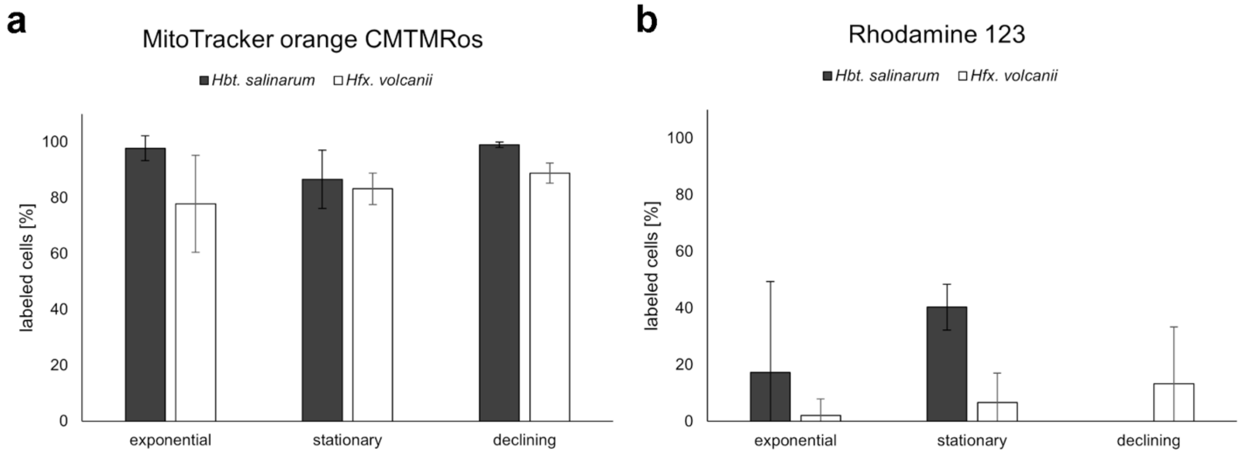


**Figure S3:** Percentage of *Hbt. salinarum* (grey) and *Hfx. volcanii* (white) labeled with either MitoTracker Orange CMTMRos or Rhodamine 123, as a function of growth phase. Cells were manually counted (biological triplicates with >10 cells each) from epifluorescence images and percentage of labeled cells was calculated as the total number of cells in fluorophore image divided by the total number of cells in correlating brightfield image. Error-bars show standard deviation of samples.

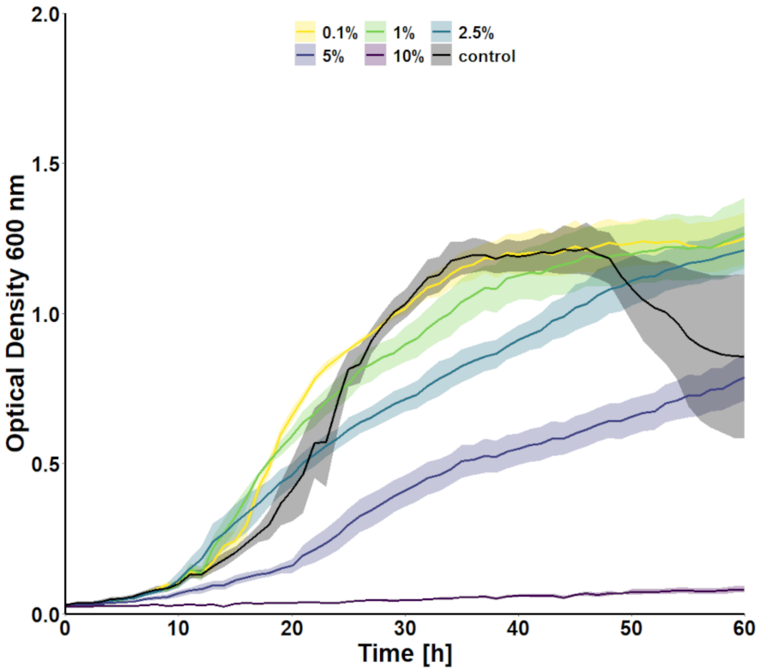


**Figure S4:** Growth impact of DMSO on *Hbt. salinarum* in complex medium. DMSO concentrations of 10 %, 5 %, 2.5 %, 1 %, 0.1 % [v/v] or without DMSO (control) were added to cultures of *Hbt. salinarum*. Growth was measured as OD600nm over shaking incubation at 37 C. Growth inhibition could be observed for concentrations of 2.5 % DMSO and higher.

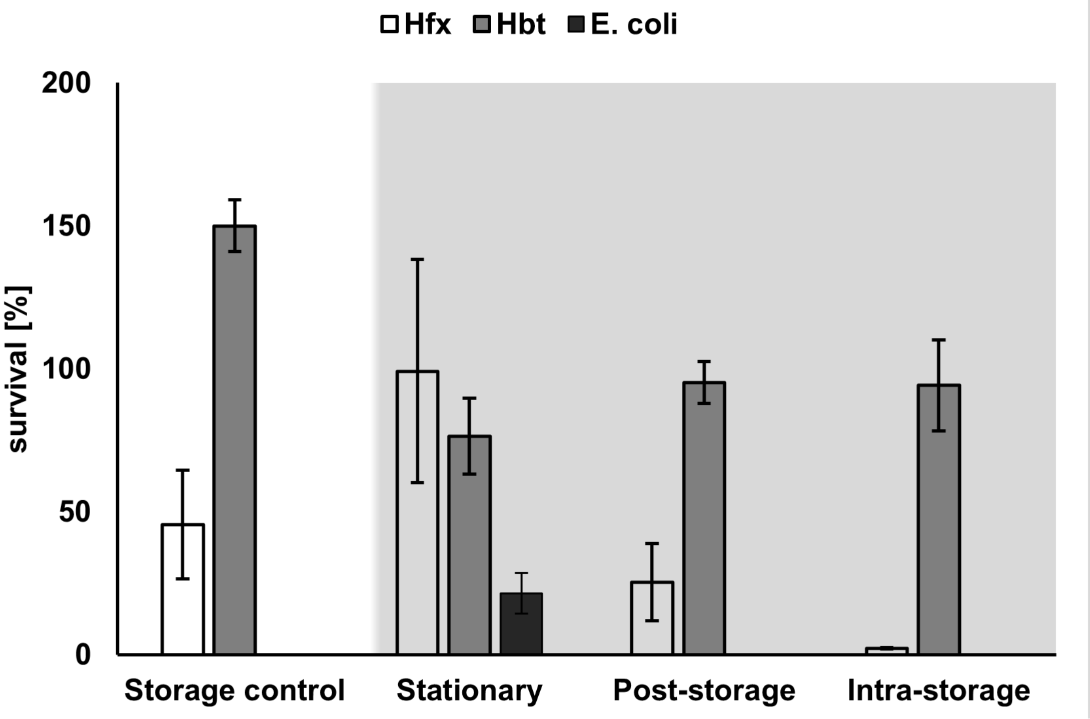


**Figure S5:** Comparison of colony forming units (CFU) for *Hbt. salinarum* (grey), *Hfx. volcanii* (clear) and *E. coli* (dark grey) with respect to both the physiological condition of the cells and the time of incubation with the LIVE/DEAD kit probes (SYTO 9, PI). Light-grey background indicating labeling experiments. Cells were either labeled immediately after reaching stationary growth phase under standard culturing conditions (37 ˚C, 180 rpm) using a 15 min incubation with both probes following the manufacturer-recommended protocol ('stationary'), or cultures were stored for an additional 5 d (room temperature, 0 rpm) reaching a decline phase to simulate entombment in halite crystals. These cells were either labelled after the secondary 5-day incubation (post-storage') or continuously exposed to SYTO 9 and PI throughout this secondary 5-day incubation ('Intra-storage') light-protected for the fluorescent probes. 'storage control' (clear background) presenting the control-incubation over 5 days. Following each treatment cells were plated on nutrient agar and incubated at 37 ˚C. Percentage survival was calculated as CFU for each incubation condition divided by CFU of ulabeled freshly stationary cultures. An increased number of CFU was observed for storage control samples of *Hbt. salinarum*, indicating cell proliferation, albeit slow, over the 5 d storage. Sample treatment scheme is shown in Figure S5.1.

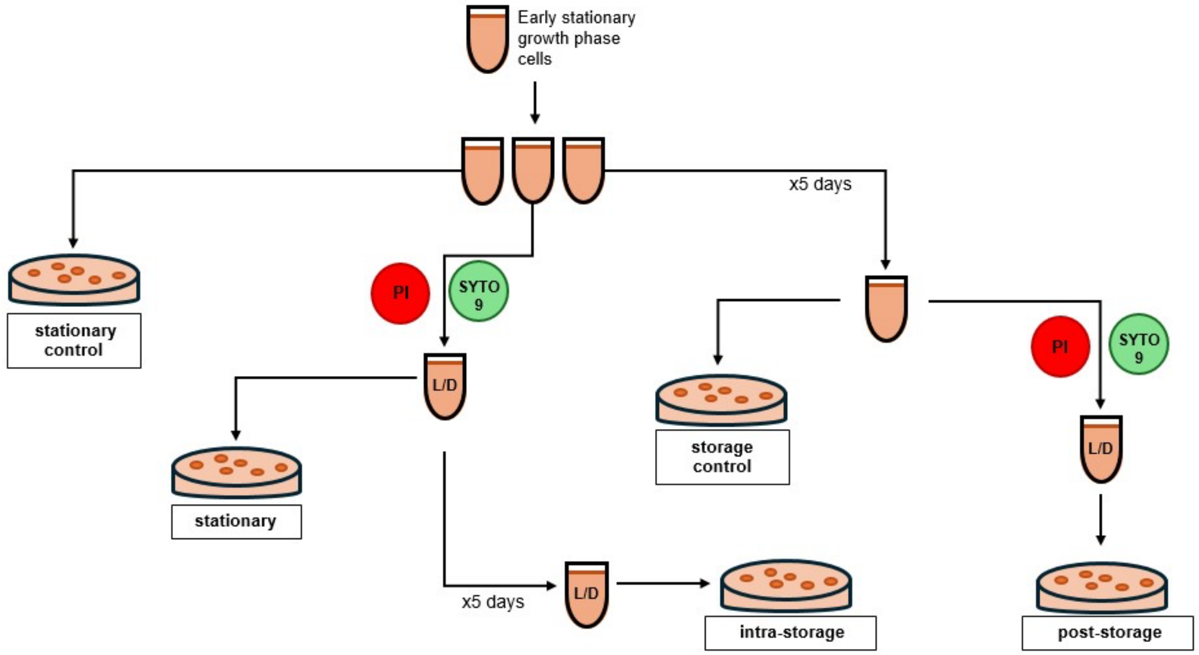


**Figure S5.1:** Method flow-chart of LIVE/DEAD kit testing with haloarchaeal cells. Early stationary growth phase cells in liquid cultures were split into three aliquots per sample. The first replicate (stationary control) was immediately plated (as serial dilution) on complex medium agar (CMA) and incubated at 37 °C to manually count colony forming units (CFU). The second replicate was labeled for 15 min at RT with Propidium Iodide (PI) and SYTO 9, the fluorescent probe components of the LIVE/DEAD kit, then either plated on CMA (stationary) or incubated over 5 days at RT under light-protection and then plated on CMA (intra-storage). The third replicate was incubated over 5 days (RT, light protected) and then either plated on CMA (storage control) or labeled for 15 min at RT with PI and SYTO 9 and then plated on CMA (post-storage).

**Table S1:** List of fluorescent probes used in this study, their biochemical function, and utilization protocols.

| | Supplier and product reference | Chemical reaction | Targeted cell envelope function | Solvent type | Stock solution | Storage temperature | Working solution | Preparation for microscopy | Optimal Ex/Em wavelengths [nm] | Ex/Em filter (POLARstar Omega® spectro-photometer) |
|---|---|---|---|---|---|---|---|---|---|---|
| **Resazurin** | Invitrogen™ (ThermoFisher™) Ref: R12204 | Reduction from blue, non-fluorescent resazurin to pink, fluorescent resorufin [-$O_2$] * | Redox activity | water | 20 mM | -20 °C | 100 µM | Incubation with 8x10E7 cells for 240 min at 37 °C, then concentrated by centrifugation (3 min, 3000 xg, RT) to 8x10E8 cells<br>Ex/Em filter cube of Nikon Epifluorescence Microscope: Ex: 560/25 Em: 607/40 | Ex: 570 Em: 580 | Ex: 544 Em: 590/10 |
| **alamarBlue™** | Invitrogen™ (ThermoFisher™) Ref: DAL1025 | Reduction from blue, non-fluorescent resazurin to pink, fluorescent resorufin [-$O_2$] * | Redox activity | Ready to use solution | 10x | 4 °C | 1x | Incubation with 8x10E7 cells for 240 min at 37 °C, then concentrated by centrifugation (3 min, 3000 xg, RT) to 8x10E8 cells<br>Ex/Em filter cube of Nikon Epifluorescence Microscope: Ex: 560/25 Em: 607/40 | Ex: 570 Em: 580 | Ex: 544 Em: 590/10 |
| **MitoTracker™ orange CMTMRos** | Invitrogen™ (ThermoFisher™) Ref: M7510 | Thiol conjugation of cysteine residues | Membrane potential | DMSO | 1 mM | -20 °C | 100 nM | Incubation with 8x10E8 cells for 60 min at 37 °C, then 2x washed with growth medium by centrifugation (3 min, 3000 xg, RT)<br>Filter cubes of Nikon Epifluorescence Microscope: Ex: 560/25 Em: 607/40 | Ex: 554 Em: 576 | Ex: 544 Em: 590/10 |
| **Rhodamine 123** | Invitrogen™ (ThermoFisher™) Ref: R302 | Always fluorescent, accumulates in cells due to its lipophilic and cationic properties | Membrane potential | DMSO | 5.25 mM | -20 °C | 15 µM | Incubation with 8x10E7 cells for 60 min at 37 °C, then 2x washed with growth medium by centrifugation (3 min, 3000 xg, RT) prior to concentration to 8x10E8 cells<br>Filter cubes of Nikon Epifluorescence Microscope: Ex: 485/20 Em: 521/30 | Ex: 507 Em: 527 | Ex: 485/12 Em: 520 |
| **Syto9 (*BacLight*™ Live/Dead™ kit)** | Invitrogen™ (ThermoFisher™) Ref: L7012 | DNA intercalation | All cells (membrane permeable probe; commonly interpreted as live cells) | Ready to use solution (DMSO) | 3.34 mM | -20 °C | 7 µM | Incubation with 8x10E8 cells for 15 min at RT<br>Laser of Zeiss LSC Microscope: Ex: 488 Em: 490/30 | Ex: 480 Em: 500 | Ex: 485/12 Em: 520 |
| **Propidium Iodide (*BacLight*™ Live/Dead™ kit)** | Invitrogen™ (ThermoFisher™) Ref: L7012 | DNA intercalation | Damaged/highly permeable cell membranes (membrane-impermeable probe; commonly interpreted as dead cells) | Ready to use solution (DMSO) | 20 mM | -20 °C | 40 µM | Incubation with 8x10E8 cells for 15 min at RT<br>Laser of Zeiss LSC Microscope: Ex: 561 Em: 561/758 | Ex: 530 Em: 625 | Ex: 530/10 Em: 620/10 |

*The resazurin-reduction can also be tracked with colorimetric methods, where resazurin-absorbance is at 600 nm and resorufin-absorbance at 570 nm.

**Supplementary Tables**

**Table S2.** Chemical composition and pH of high-salt solutions for *Hbt. salinarum* and *Hfx. volcanii* based on their respective complex growth media CM/CM+ and HvYPC. *Solutions contained either only the salts needed for each organism (HfxS and HbtS) or the salts plus only one of the main complex organic nutrient sources: Oxoid® neutralized peptone (Pep N), Oxoid® non-neutralized peptone (Pep NN) Bacto™ yeast extract (YE) and Bacto™ Casamino acids (Cas).*

| | HbtS | HbtS PepN | HbtS PepNN | CM | CM+ | HfxS | HfxS PepN | HfxS PepNN | HfxS YE | HfxS Cas | HvYPC |
|---|---|---|---|---|---|---|---|---|---|---|---|
| pH | 6.31 | 6.32 | 5.86 | 7.2 | 7.1 | 6.85 | 7.02 | 6.99 | 6.44 | 6.69 | 7.1 |
| NaCl [M] | 4.28 | 4.28 | 4.28 | 4.28 | 4.28 | 2.46 | 2.46 | 2.46 | 2.46 | 2.46 | 2.46 |
| MgSO4 · 7 H2O [mM] | 81.2 | 81.2 | 81.2 | 81.2 | 81.2 | 85.2 | 85.2 | 85.2 | 85.2 | 85.2 | 85.2 |
| KCl [mM] | 26.8 | 26.8 | 26.8 | 26.8 | 26.8 | 56.3 | 56.3 | 56.3 | 56.3 | 56.3 | 56.3 |
| C6H5Na3O7 · 2 H2O [mM] | 10.2 | 10.2 | 10.2 | 10.2 | 10.2 | | | | | | |
| MgCl2 · 6 H2O [mM] | | | | | | 88.5 | 88.5 | 88.5 | 88.5 | 88.5 | 88.5 |
| Non-Neutralized Peptone (Oxoid® REF LP0037) | | | 1 % w/V | | | | | 0.1 % w/V | | | |
| Neutralized Peptone (Oxoid® REF LP0034) | | 1 % w/V | | 1 % w/V | | | 0.1 % w/V | | | | 0.1 % w/V |
| Bacto™ Yeast Extract (gibco, REF 12750) | | | | | | | | | 0.5 % w/V | | 0.5 % w/V |
| Bacto™ Casamino acids (gibco, REF 223050) | | | | | | | | | | 0.1 % w/V | 0.1 % w/V |
| Tris HCl (pH 7.5) [mM] | | | | | | | | | | | 12 |
| Glycerol | | | | | 0.50% | | | | | | |
| ZnSO4 · 7 H2O [mM] | 5.97 | 5.97 | 5.97 | | 5.97 | | | | | | |
| MnSO4 [µM] | 21.9 | 21.9 | 21.9 | | 21.9 | | | | | | |
| CuSO4 · 7 H2O [µM] | 2.73 | 2.73 | 2.73 | | 2.73 | | | | | | |
| FeSO4 [mM] | 23.03 | 23.03 | 23.03 | | 23.03 | | | | | | |
